# MTB-KB: A Curated Knowledgebase of *Mycobacterium tuberculosis* Related Studies

**DOI:** 10.64898/2026.04.07.716833

**Authors:** Pan Li, Cuidan Li, Rongxi Zhu, Wenjing Sun, Hengyu Zhou, Zhuojing Fan, Liya Yue, Sijia Zhang, Xiaoyuan Jiang, Quan Luo, Jinying Han, Hairong Huang, Adong Shen, Tuohetaerbaike Bahetibieke, Jing Wang, Wenbao Zhang, Hao Wen, Haitao Niu, Congfan Bu, Zhang Zhang, Jingfa Xiao, Renjun Gao, Fei Chen

## Abstract

Tuberculosis (TB), caused by *Mycobacterium tuberculosis* (MTB), has regained its position as the world’s leading killer among infectious diseases. Despite extensive research progress across epidemiology, diagnosis, drug development, treatment regimens, vaccines, drug resistance, virulence factors, and immune mechanisms, MTB-related knowledge remains fragmented across thousands of publications, limiting its effective use. To address this gap, we present MTB-KB, a literature-curated knowledgebase that systematically integrates high-impact findings from eight major sections of TB research. The current release contains 75,170 associations from 1,246 publications, covering 18,439 entities standardized using authoritative databases and WHO-endorsed classifications. A central feature is the interactive knowledge graph, which links cross-section associations to reveal and infer MTB–host interactions, treatment strategies, and vaccine development opportunities. MTB-KB also provides a user-friendly interface with browsing, advanced search, and statistical visualization. Overall, by consolidating dispersed MTB knowledge into a structured and accessible platform, MTB-KB provides a valuable resource for researchers, clinicians, and policymakers, supporting both basic and clinical TB research, enabling evidence-based TB prevention, diagnosis, and treatment, and contributing to global elimination efforts. MTB-KB is accessible at https://ngdc.cncb.ac.cn/mtbkb/.

## Introduction

Tuberculosis (TB), caused by *Mycobacterium tuberculosis* (MTB), regained its position in 2023 as the world’s leading cause of death from a single infectious agent. According to the World Health Organization (WHO), there were about 10.7 million new TB cases in 2024, and an estimated 1.23 million people died from the disease [1]. Despite major advances in TB diagnosis [2, 3], treatment [4], and prevention [5] over the past century, the WHO’s End TB target by 2035 remains far from being achieved, and the disease continues to pose a serious threat to global public health.

Over recent decades, research has greatly advanced our understanding of TB across multiple fields, which has significantly contributed to TB prevention, control, and treatment. In epidemiology, the temporal and spatial distribution, drug resistance patterns, and other characteristics of the eight main MTB lineages have been extensively studied [6]. In diagnostics, molecular tools such as Xpert MTB/RIF Ultra [7] have improved the speed and accuracy of detecting TB and drug resistance. New drugs, including delamanid [8], linezolid [9], bedaquiline [10], and pretomanid [11], have improved treatment outcomes for patients with drug-resistant TB. Numerous new resistance mutations and mechanisms have been elucidated; for example, approximately 95% of rifampicin resistance mutations occur within an 81-bp core region of *rpoB* [12]. Virulence studies have identified many novel virulence factors and infection mechanisms, such as the ESX secretion system [13]. Host immunology research has revealed the critical role of interferon-gamma (IFN-γ) and Th1 immune responses in protection [14] against MTB, with key genes including *IFNG*, *IFNA*, and *STAT1* [15]. Several vaccine candidates, such as M72/AS01E, have shown encouraging results in phase II clinical trials [16].

However, these findings are scattered across thousands of publications, and without systematic integration, valuable insights are often overlooked or underutilized. Systematic integration of these information into a unified and well-curated MTB knowledgebase can offer valuable references for TB prevention, diagnosis and treatment, empower researchers, clinicians, and policymakers, and provide valuable support toward achieving the WHO’s goal of ending the global TB epidemic by 2035.

To date, several MTB-related databases have been created, each serving specific research needs. MycoBrowser [17] provides genome annotations for *Mycobacterium* species. GMTV [18] catalogs single nucleotide polymorphisms (SNPs) and insertion and deletion (indels) from over 1,000 MTB strains, along with drug resistance, clinical stage, and geographic data. BioCyc [19] offers curated biological pathways and omics tools, while MTBSD [20] focuses on MTB protein structures to support structure-based drug design. SITVITWEB [21] compiles genotyping data from more than 62,000 clinical isolates in 153 countries, supporting lineage classification through spoligotyping, MIRU-VNTRs, and ETR. TB Portals [22] integrates case-level data from drug-resistant TB patients, combining clinical records, radiological images, and genomic sequences, with built-in analytical and visualization tools.

While these databases provide indispensable genomic, clinical, or functional datasets, they are mostly data-driven and rely on raw or preprocessed datasets, and specialized in a single data type or research section. Therefore, a curated knowledgebase that organizes key findings from the literature into a structured and accessible resource could better support TB research.

To address these issues, we present MTB-KB (https://ngdc.cncb.ac.cn/mtbkb/) to complement these data-driven repositories by providing a curated knowledgebase that organizes high-impact findings across eight major sections of tuberculosis research: *Epidemiology, Diagnosis, Drug, Regimen, Vaccine, Drug Resistance, Virulence Factor, and Immune Mechanism*. MTB-KB employs a rigorous manual curation process to extract key associations from highly cited publications. Thousands of curated entities are further enriched and standardized using authoritative external databases, ensuring semantic consistency, interoperability, and broader knowledge integration. Importantly, MTB-KB offers an interactive knowledge graph, enabling users to explore cross-sectional knowledge association and inference. As a publicly accessible resource, MTB-KB integrates a broad spectrum of MTB knowledge into a unified, user-friendly platform for exploring tuberculosis-related knowledge and supporting further research in TB biology, diagnosis, and treatment. and contributing to global efforts in TB control, treatment, and prevention.

## Data curation and database development

### Knowledge curation and integration

To ensure the quality and uniformity of data in MTB-KB, we established a three-step knowledge curation workflow: literature search and filtering, manual study curation, and association collection and integration.

*Literature search and high-impact publication filtering:* To comprehensively capture diverse aspects of MTB research, MTB-KB content was organized into eight core sections: *Epidemiology, Diagnosis, Drug, Regimen, Vaccine, Drug Resistance, Virulence Factor, and Immune Mechanism*. This framework reflects the main research directions of MTB studies and enables systematic curation from the macro to the micro level, spanning molecular, clinical, and translational fields.

A comprehensive search was conducted in the Web of Science Core Collection using predefined, section-specific keyword combinations to ensure targeted retrieval across MTB-KB’s eight core research sections (Supplementary table S1). For example, the Diagnosis section used queries such as “Diagnosis,” “Diagnostic,” and “Identification” combined with “Tuberculosis”. Similarly, tailored keywords were used for other sections, including *Epidemiology* (“Outbreak”, “Epidemic”, “Prevalent”), *Treatment* (“Treatment”, “Therapy”), *Vaccine* (“Vaccine”), *Virulence Factor* (“Virulence”, “Virulence factor”), *Drug Resistance* (“drug resistance”, “drug susceptibility”, “antibiotic resistance”, “drug sensitivity”), and *Immune Mechanism* (“Immune”). Notably, the *Drug* and *Regimen* sections were initially retrieved under a unified treatment-related query, then separated during curation based on distinct reporting patterns.

This process ensured a highly sensitive retrieval of MTB-relevant literature, resulting in a total of 166,698 initial publications. To establish a high-quality, manually verified evidence foundation, we prioritized these publications using the normalized indices (z-index, total citations divided by year since publication) [23] to identify high-impact studies. Unlike raw citation counts, the z-index normalizes for both time and research domain. We further selected the top 5% within each section to ensure that the initial core of the knowledgebase reflects the most widely recognized and validated findings in the field. Finally, we identified 8,227 highly cited publications for subsequent manual review, as detailed in our PRISMA-style workflow (Supplementary Figure 1).

*Manual literature curation:* The filtered publications were manually reviewed to retain only those reporting experimental findings, methodologies, or biological insights directly relevant to the predefined curation schema of MTB-KB. For each research section, we define the key entities, their relationships, and the relevant contextual information to be recorded. To ensure standardization, all entries are normalized using controlled vocabularies and linked to its source publication to preserve traceability and context (Supplementary Table S2).

For example, in the *Immune Mechanism* section, the initial keyword and z-score based filtering yielded 1,005 publications. We first removed non-primary research articles, including reviews (n = 50), meta-analyses (n = 33), guidelines (n = 10), case studies (n = 10), and letters (n = 2). Subsequent manual screening of abstracts and relevant keywords excluded studies outside the scope of host immune mechanism, such as those focusing on clinical trials or vaccine development (n = 177), bacterial metabolism (n = 43), or gut microbiota (n = 7), resulting in 563 candidate publications. Each remaining study was then assessed against the immune-related curation model, which records three dimensions of immune response at cell, antibody, and gene/cytokine/chemokine levels. For each level, we recorded the detailed information including the affected entity, up/down-regulation, experimental methods, and statistical measures (*e.g.*, p-value, fold change), along with shared metadata such as experimental species, case/control definitions, sample types, and organism. Applying these criteria, 73 publications were ultimately retained.

The same process with section-specific curation models was applied to all the sections, yielding the following retained publication counts: *Epidemiology* (n=117), *Diagnosis* (n = 334), *Drug* (n = 194), *Regimen* (n = 247), *Vaccine* (n = 134), *Drug Resistance* (n = 210), and *Virulence Factor* (n = 78). In total, 1,246 publications were ultimately curated across all sections. For each included study, we recorded core metadata including bibliographic details (*e.g.*, authors, journal, title), data sources (*e.g.*, GSE or NCT accession numbers), and section-specific findings.

*Associations Collection and Integration:* From each curated publication, we manually extracted detailed associations for the relevant section(s). Four inclusion criteria were applied: (i) statistically significant results (typically p < 0.05), (ii) validated mechanisms of action, (iii) demonstrated clinical efficacy (*e.g.*, treatment success, culture conversion, or time to sputum negativity) or experimental efficacy (*e.g.*, MIC changes, CFU reduction), and (iv) molecular or genetic linkages. To ensure consistency, each association in MTB-KB is centered on a key entity and includes supporting details such as experimental methods, related entities, and literature sources.

In addition to curated associations, MTB-KB provides structured basic information on MTB, including general descriptions, taxonomic classification, and curated lists of related diseases and associated clinical symptoms. Disease and symptom annotations are based on the Disease Ontology (DO) [24], supplemented by information from DO records and relevant literature. This layer gives users quick access to essential background information for both navigation and contextual understanding.

Finally, all entity names were standardized to ensure consistency across the knowledgebase. This normalization step allows relationships between sections to be harmonized, enabling unified searches and network-based analyses.

### Entity annotation and classification standards

To ensure semantic consistency and interoperability, all curated entities in MTB-KB are annotated using authoritative external resources. Diseases are mapped to the Disease Ontology (DO) [24], and their associated signs and symptoms follow the Human Phenotype Ontology (HPO) [25]. Drug annotations are sourced from DrugBank [26], while vaccine annotations use both DrugBank and VIOLIN [27]. MTB genes are aligned with entries from MycoBrowser (17), and virulence factors are described according to the Virulence Factors Database (VFDB) [28]. Host immune genes are mapped to Ensembl [29] and HGNC [30], while immune cells are standardized using Cell Taxonomy [31]. Where applicable, synonyms and aliases are included to support flexible searches and improve user navigation.

To enhance clinical relevance, MTB-KB integrates WHO-endorsed classification frameworks. Anti-TB drugs are categorized according to the *WHO Operational Handbook on Tuberculosis* [32], which groups medicines by efficacy, safety, and role in treatment regimens. Resistance-associated genes and mutations follow the *WHO Catalogue of Mutations in Mycobacterium tuberculosis Complex and Their Association with Drug Resistance* (2nd edition) [33], which also assigns confidence levels to genotype–phenotype associations.

### Knowledge graph construction

The interactive knowledge graph is publicly accessible through the MTB-KB web interface (https://ngdc.cncb.ac.cn/mtbkb/KnowledgeGraph). The MTB-KB knowledge graph was designed to systematically organize and visualize literature-curated associations across key sections of the database. It integrates five primary entity types: drugs, vaccines, virulence factors, MTB genes, and host immune genes. To ensure consistency and interoperability, we have clarified that curated entities are normalized using standardized names from authoritative resources: drug annotations are sourced from DrugBank, vaccine annotations use both DrugBank and VIOLIN, MTB genes are aligned with entries from MycoBrowser, while virulence factors are described according to the Virulence Factors Database, host immune genes are mapped to Ensembl and HGNC. Each association is classified into one of four interaction types: regulation, functional impact, targeting, or containment (Supplementary Figure 2). To improve interpretability, each relationship is annotated with the number of supporting evidence entries. This value represents the frequency of reported evidence for a given association in the literature and is used as the edge weight in the graph. For visualization, the node size reflects connectivity and the edge weights represent the frequency of literature evidence. The graph uses a dynamic force-directed layout, allowing users to adjust the view in real time. The visualization interface is implemented using Apache ECharts, enabling interactive exploration directly within the web browser.

To keep complex visualizations clear, MTB-KB employs a smart node selection strategy [34]. For a given central node and exploration depth (1–3 degrees), up to 50 high-relevance nodes are prioritized based on: (i) connection strength, (ii) evidence support, and (iii) diversity of entity types. Users can also filter nodes through the graph legend, controlling which entity types are displayed to focus on specific research needs. The graph is fully interactive, supporting hovering, clicking, dragging, and re-centering, enabling flexible, multi-angle exploration of curated MTB knowledge.

### Database implementation

MTB-KB was developed using Django (v4.2.23, https://www.djangoproject.com/) as the backend framework and Vue 3 (https://vuejs.org/) for the frontend. MySQL (v8.0.42, https://www.mysql.com/) serves as the database engine for structured data storage and querying. ECharts (https://echarts.apache.org/) is used for rendering interactive statistical charts, and Bootstrap Icons (https://icons.getbootstrap.com/) support lightweight user interface elements.

## Database contents and usage

MTB-KB provides a comprehensive, literature-curated integration of *Mycobacterium tuberculosis* and tuberculosis knowledge across eight major research sections (Figure 1). All content is manually extracted from highly cited publications and structured into standardized formats to ensure accuracy and ease of interpretation. Curated entities are annotated using authoritative external databases and WHO-endorsed classifications. In addition to tables, MTB-KB offers interactive features including search, summary statistics, and a dynamic knowledge graph, allowing users to explore complex relationships across sections. By systematically integrating scattered research, MTB-KB makes it easier to apply existing knowledge to future studies and public health efforts.

**Figure 1.**
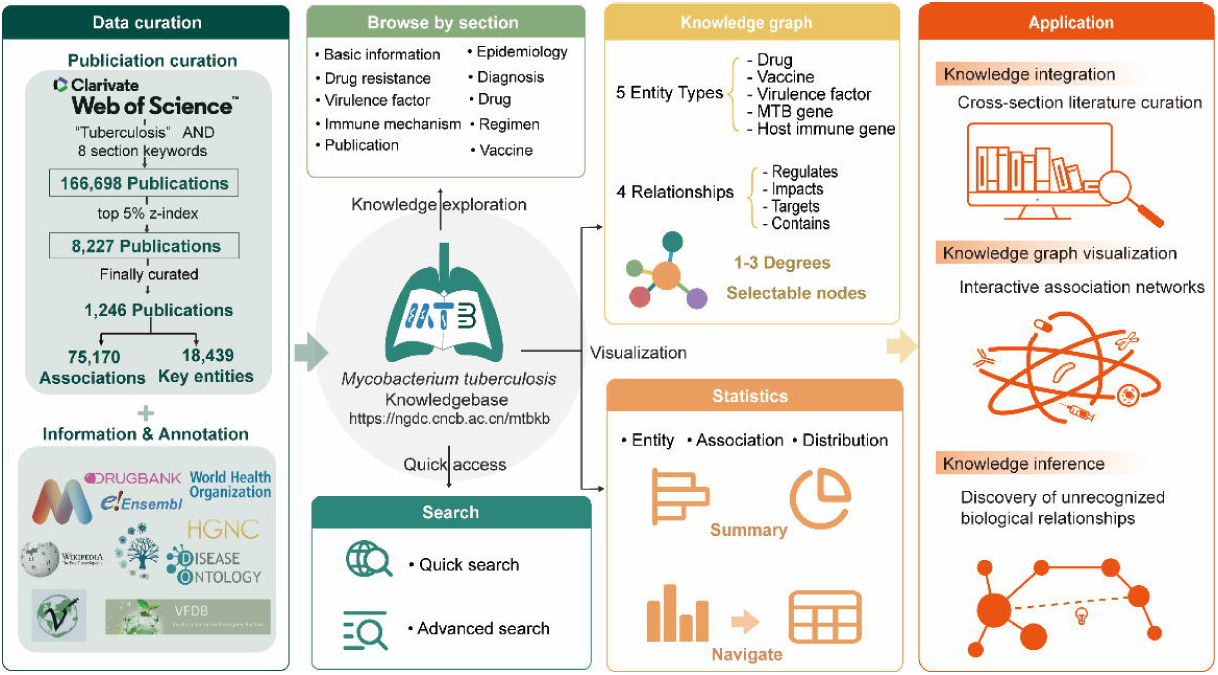
Overview of MTB-KB curation workflow and modules The figure summarizes the data curation workflow, organization of curated content, and key functional modules of MTB-KB. Associations are curated from published studies and organized into eight sections, enabling structured browsing, search, and statistical overview. An interactive knowledge graph visualizes evidence-linked associations among 5 major entity types and supports multi-degree exploration. The Application panel highlights the main ways MTB-KB can be used, including integration of knowledge across research sections, visualization of curated associations, support for knowledge inference.

### Comprehensive association knowledge for MTB-KB

The current release covers curated associations across eight sections: *Epidemiology, Diagnosis, Drug, Regimen, Vaccine, Drug Resistance, Virulence Factor, and Immune Mechanism*. A total of 75,170 literature-supported associations were extracted from 1,246 high-impact publications (Figure 2A), involving 18,439 unique biological entities: 584 to pathogen features, 16,673 related to host immune responses, and 1,182 to clinical applications (Figure 2B). All associations link to their source publication and are annotated with study metadata, experimental context, and clinical relevance.

**Figure 2.**
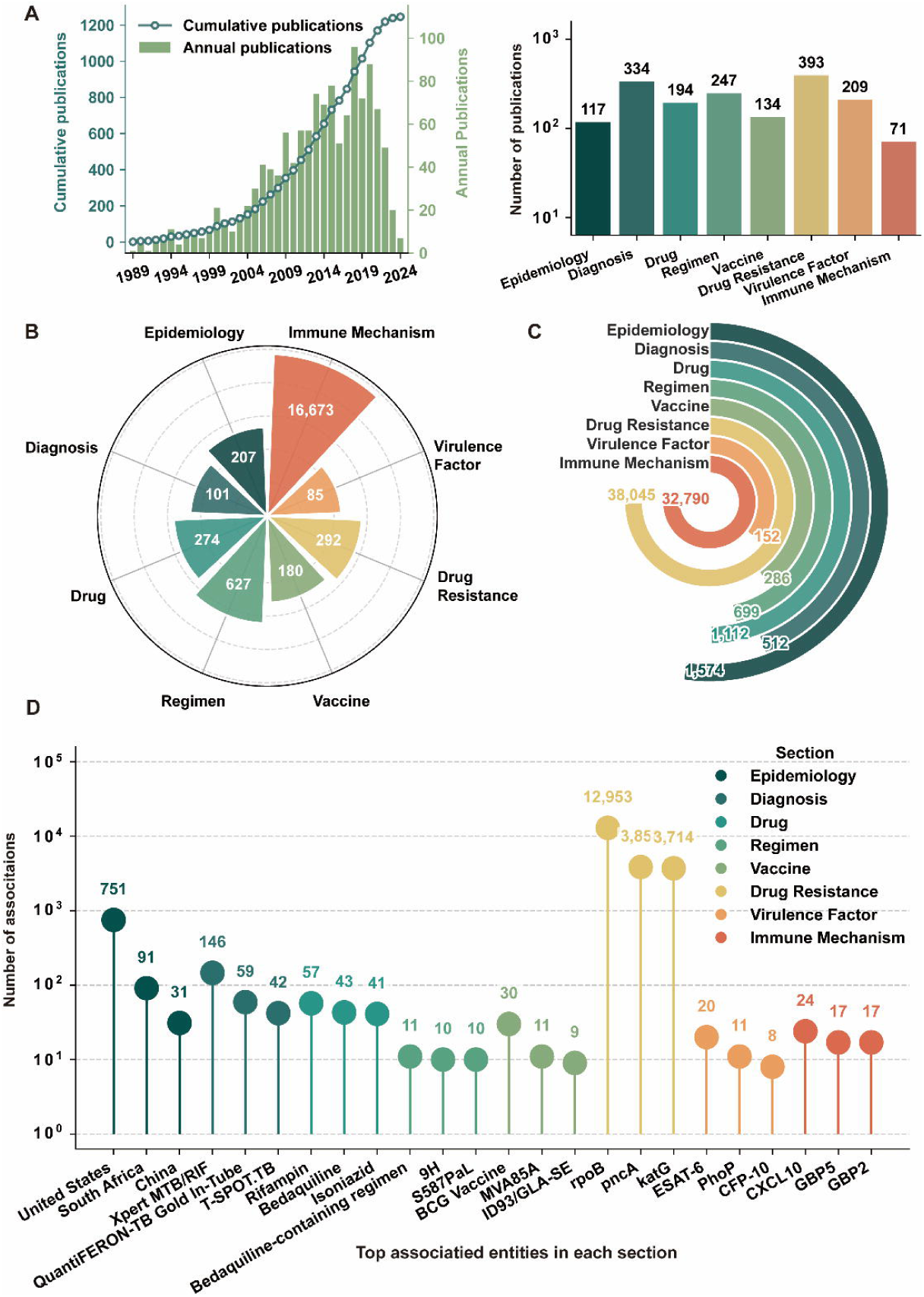
Statistics derived from curated MTB-related entities and associations **A.** Distribution of the 1,246 curated publications by year (left) and by MTB research section (right). **B.** Distribution of 18,439 unique curated biological entities across eight MTB research sections. **C.** Distribution of 75,170 literature-supported associations across research sections. **D.** Top three most frequently associated entities in each research section.

Pathogen-specific entities include 292 drug resistance genes and 85 virulence factors. For example, the drug resistance genes, *rpoB*, *katG*, and *inhA* are reported in 108, 100, and 74 curated publications, respectively. For virulence factors, the knowledgebase captures well-characterized factors such as phthiocerol dimycocerosates (PDIM) and CFP-10, whose critical role in MTB pathogenicity are well documented.

Host-derived genes, ‘make up the majority (16,594; 90%) of curated entities. For example, *CXCL10*, reported in 14 publications, is highlighted as both a protective factor and a potential diagnostic marker [35]. MTB-KB also includes 72 immune cell types (*e.g*., plasmacytoid dendritic cell) and 7 antibodies (*e.g*., anti-Ag85B antibody), enabling detailed study of cellular and antibody-mediated interactions.

Clinical application-related entities include 274 drugs (*e.g*., bedaquiline, rifampin), 627 treatment regimens (*e.g.*, 2HREZ/4HR, BPaMZ), 180 vaccines (*e.g*., BCG, MVA85A), and 101 diagnostic methods (*e.g.*, Xpert MTB/RIF, QuantiFERON-TB

Gold Plus). Many entities appear in multiple roles: IL-1β participates in immune responses and is studied as a drug target; CD4+ T cells are involved in both natural immunity and vaccine responses; and several resistance genes (*e.g*., *rpoB, inhA*) serve as both drug targets and diagnostic markers. This interconnected landscape supports multi-dimensional exploration of TB biology and clinical interventions.

The 75,170 associations form a structured network of experimentally supported relationships, dominated by drug resistance (38,045; 51%) and immune mechanisms (32,790; 44%) (Figure 2C). Figure 2D highlights the top three most-associated entities in each section. In the *Diagnosis* section, Xpert MTB/RIF leads with 146 associations, underscoring its role in rapid TB detection and resistance testing. In *Drug*, rifampin (57 associations) and bedaquiline (43 associations) are most prominent (rifampin as a long-standing first-line therapy and bedaquiline as a newer multidrug-resistant tuberculosis (MDR-TB) drug [10, 36]). In *Immune Mechanism*, guanylate-binding proteins *GBP5* and *GBP2* each have 17 associations, reflecting growing interest in innate immunity pathways for TB control [37]. Together, these curated associations provide a comprehensive view of MTB-related relationships and make the knowledge readily accessible for diverse research and application needs.

### User-friendly modules for data access

*Browse:* Provides section-specific views for all eight research areas: *Epidemiology, Diagnosis, Drug, Regimen, Vaccine, Drug Resistance, Virulence Factor, and Immune Mechanism* (Figure 3A). The section contains a summary table listing key entities, including key attributes such as name, category, and number of associated records. Each entity links to a detailed page showing basic information and all related associations. A Basic Information page offers background on MTB, including related diseases and clinical symptoms sourced from the Disease Ontology. A separate publication view provides citation details (*e.g.*, title, journal, year, PMID, DOI) and organizes related entries by research section.

**Figure 3.**
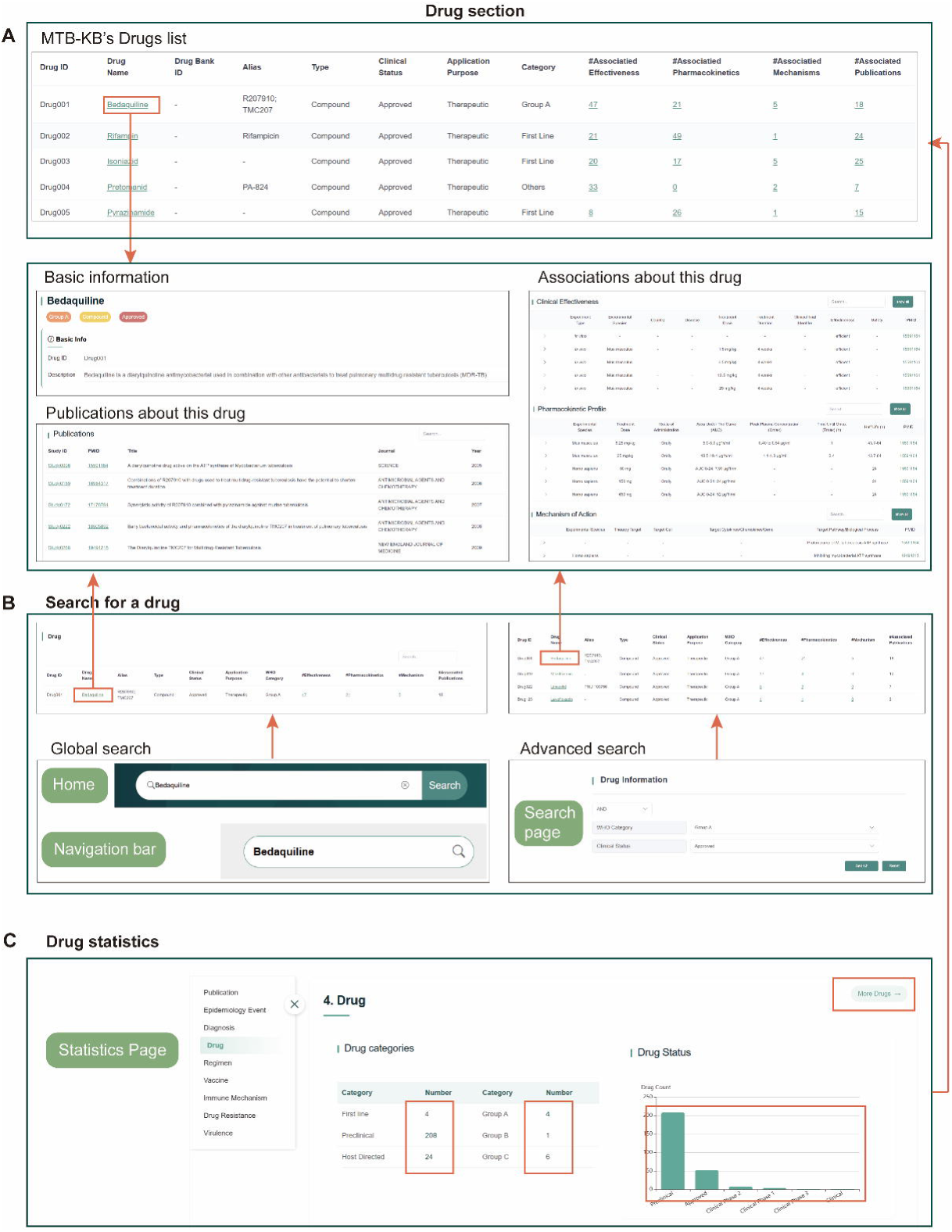
User interfaces of MTB-KB’s Drug Section for browsing, searching, and statistical analysis **A.** Browse module for drugs, showing a summary table with key attributes (drug name, aliases, category, targets, associated resistance genes, and linked publications) and a dedicated detail page with basic information, relevant associations, and related publications. **B.** Search module, including global search (left) and advanced search (right), enabling queries of drug-related associations by entity names or keywords. **C.** Statistics module for drugs, providing real-time analytics of drug categories and status.

*Search:* Offers quick keyword search across all sections and advanced search in three categories: Prevention, Treatment and Diagnosis, Host Information, and MTB Information (Figure 3B). For advanced search, filters allow precise queries, such as by gene name, cytokine type, immune cell, MTB gene, or virulence factor. Results are presented in structured tables with direct links to entities and publications.

*Statistics:* Provides an interactive visual overview of curated content across all eight research sections (Figure 3C). It summarizes entities, associations, and publications through charts and tables, with dedicated panels for each section showing key indicators such as the number of curated entities, literature-supported associations, and publication counts. Charts are organized by section, typically offering 2–3 statistical views tailored to its content. For example, in *Drug Resistance*, panels display top-associated drugs, numbers of resistance mutations, and resistance categories; in *Immune Mechanism*, word clouds visualize the frequency of immune components such as genes or cells. Publication statistics, covering yearly trends and journal distribution, are provided separately. All visualizations are linked to underlying data tables for direct access, enabling smooth navigation from summaries to detailed records.

*Download:* Provides public access to all curated data, including publication lists, entity annotations, and clinical trial information, ensuring transparency, reusability, and ease of offline analysis.

### Systematically integrated knowledge graph

MTB-KB integrates associations among drugs, vaccines, MTB virulence factors, MTB genes, and host immune genes into a comprehensive relationship network presented as an interactive knowledge graph. This graph serves as a powerful tool to uncover biological networks underlying MTB-host interactions, informing treatment strategies, and guiding vaccine development.

The graph includes five node types (101 drugs, 158 vaccines, 85 virulence factors, 640 MTB genes, and 505 host immune genes), four interaction types (770 impacts, 751 targets, 363 regulates, and 256 contains) and supports adjustable exploration depths (1–3 degrees). Visual encodings improve interpretability: node size represents connectivity, and edge thickness reflects the strength of literature support. Users can filter by entity type, rearrange nodes via drag-and-drop, and export high-resolution images for publication or presentation. These customization features allow targeted, flexible exploration tailored to specific research interests.

Case study – drug resistance: The utility of the knowledge graph is exemplified by its analysis of drug resistance patterns (Figure 4A). Among anti-TB drugs, isoniazid forms the largest network, connecting to 118 entities and supported by 5,850 associations from 110 publications. It shows strong links to *katG* (3,676 associations), *fabG1* (534 associations), *inhA* (423 associations), *mshA* (430 associations), and *ahpC* (187 associations)—all classified as Tier 1 resistance determinants in the WHO Catalogue [33]. Rifampin ranks second, associated with 72 entities through 14,763 associations from 110 publications. Its strongest connection, highlighted in the graph by the thickest edge, is with *rpoB* (12,707 associations), the primary drug target [38], followed by *rpoC* (1,656 associations) and *rpoA* (165 associations). These findings are consistent with the central clinical roles of isoniazid and rifampin, which are the most widely used anti-TB drugs, and therefore have been the focus of extensive and in-depth drug resistance research [39, 40].

**Figure 4.**
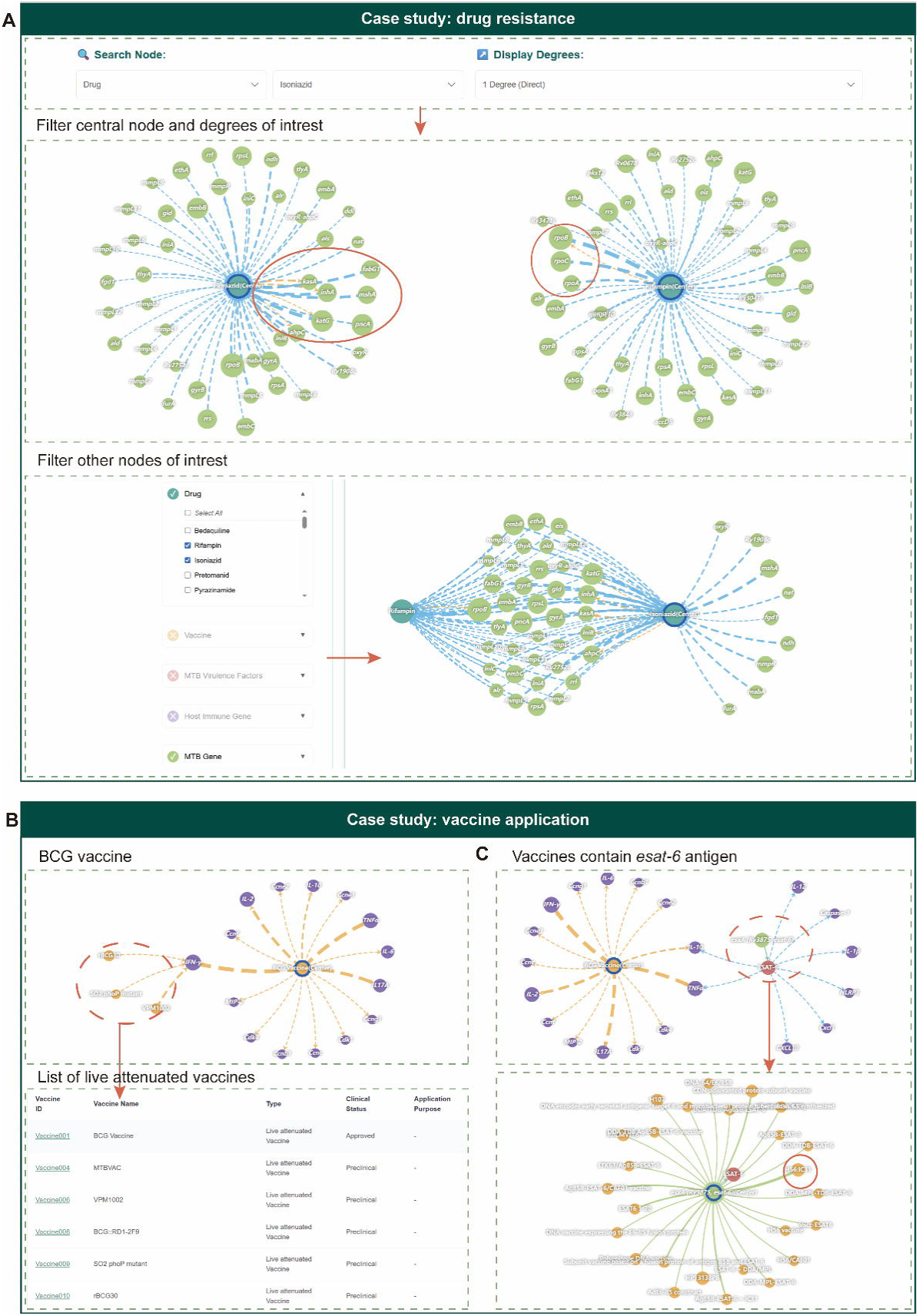
Application examples of MTB-KB’s interactive knowledge graph **A.** The top panel shows the central-node selection interface. Drug resistance case study: Isoniazid-centered (left) and rifampin-centered (right) networks, with shared resistance-associated entities highlighted. The bottom panel shows filtered views focusing on drug resistance genes using node-type selection. **B.** Vaccine case study – BCG: BCG-centered immune network highlighting strong associations with IFN-γ (top left) and live attenuated vaccines (top right); the bottom panel lists curated live attenuated vaccine candidates. **C.** Vaccine case study – ESAT-6: BCG-centered network (top) showing no direct link to ESAT-6 due to RD1 deletion; ESAT-6-centered network (bottom) highlighting associated vaccine candidates, including H56:IC31.

Case study – vaccine application: Beyond integrating and visualizing known associations, MTB-KB’s knowledge graph supports the inference of indirect relationships. For example, the BCG-centered subnetwork positions BCG as a key hub linking multiple host immune components (Figure 4B). Its strongest connection is with IFN-γ, supported by six curated studies. The graph also highlights links between BCG and recombinant BCG (rBCG) strains engineered to enhance IFN-γ responses [41]. Importantly, IFN-γ is not only central to anti-tuberculosis immunity but also functions as a critical effector in cancer immunotherapy. Consequently, rBCG strains originally designed for TB prevention may have unexpected translational potential.

As previously reported, intravesical BCG is a cornerstone immunotherapy for non-muscle-invasive bladder cancer (NMIBC) [42], but its efficacy requires frequent instillations, often causing local and systemic adverse effects [43]. The knowledge graph suggests a mechanistic convergence: rBCG strains that sustain IFN-γ induction for TB vaccination could be repurposed to reduce instillation frequency while maintaining antitumor efficacy, potentially improving treatment tolerability in NMIBC [44]. This repurposing opportunity, identified through network-based reasoning in MTB-KB, illustrates how pathogen-targeted engineering can inform solutions for clinically distinct yet mechanistically related conditions.

In parallel, the graph also helps bridge important gaps. For example, despite BCG is a hot hub in the immune network, it lacks a direct connection to the critical virulence factor ESAT-6 (Figure 4C), due to the deletion of the RD1 region (including ESAT-6) in the BCG genome [45, 46]. This absence results in insufficient stimulation of ESAT-6–specific T cell responses—a known limitation of BCG in conferring long-term protection [47]. The network pattern points to a compensatory strategy: supplementing BCG vaccination with ESAT-6–containing subunit vaccines to restore the missing immunogenic components.

This insight is reinforced by the ESAT-6–centered graph, which highlights multiple vaccine candidates. Among them, H56:IC31 is prominently connected. H56 (comprising Ag85B, ESAT-6, and Rv2660c) has been shown to boost Th1 responses in BCG-primed non-human primates [48], and phase I/IIa trials have demonstrated promising safety [49] and immunogenicity [50] in endemic populations. These network-derived insights support the rational design of complementary vaccination strategies, such as BCG priming followed by ESAT-6–based boosting, facilitating mechanism-guided TB vaccine development.

Collectively, the knowledge graph in MTB-KB integrates knowledge across sections, from MTB drug resistance/virulence to immune mechanisms, vaccines, and treatments. The graph not only visualizes existing knowledge but also facilitates hypothesis generation and multi-dimensional analysis, enabling researchers to identify novel patterns and explore previously unrecognized biological relationships. By bridging foundational research and clinical application, MTB-KB offers a valuable platform for advancing TB research, control and elimination.

## Discussion and future developments

MTB-KB provides a unified, literature-curated knowledgebase that systematically organizes high-impact findings across eight key sections, covering the major research axes of MTB-related studies. Unlike existing databases primarily compile raw datasets, MTB-KB features three key innovations:

Manual curation of high-impact literature: MTB-KB systematically extracts associations from carefully selected, highly cited publications, with an emphasis on cross-domain integration, connects host immune responses, pathogen characteristics, and clinical interventions within a single structured platform.

Standardized annotation and ontology integration: All curated entities are enriched with additional information and further standardized using authoritative external ontologies and classifications, thereby improving consistency, interoperability, and clinical relevance.

Interactive knowledge graph construction: Beyond static listings, MTB-KB offers an interactive knowledge graph that systematically organizes curated associations, enabling users to intuitively explore multi-level relationships and derive new insights from existing knowledge. Through this integration and visualization, MTB-KB moves beyond a static data list to become a practical platform for multi-dimensional research exploration and evidence-based clinical strategies that contribute to TB elimination efforts.

TB, as an ancient yet persistent threat to human health, has generated an extensive body of literature. With the rapid emergence and advancement of cutting-edge biotechnologies (such as high-throughput sequencing and artificial intelligence), the volume of TB-related publications will continue to grow. As the first and currently the only dedicated MTB knowledgebase, MTB-KB has taken an important step in the collection, integration, sharing, and utilization of vast TB literature, serving as a valuable resource for both basic and clinical TB research and facilitating TB prevention, diagnosis, and treatment. Currently, the knowledgebase curates a targeted subset of influential literature, prioritizing studies with higher citation-normalized indices (z-index) to establish a high-quality, manually verified evidence foundation. While the z-index mitigates time-related bias by normalizing citations across publication years, we acknowledge that citation-based metrics may still underrepresent novel, recent studies with lower initial counts. In addition, given the enormous volume of MTB-related literature, relying solely on manual curation makes it difficult to achieve comprehensive coverage of all relevant research domains (from fundamental biology to translational and clinical studies).

In the future, as a core resource of the National Genomics Data Center (NGDC, https://ngdc.cncb.ac.cn), MTB-KB is committed to continuous, long-term updates on an annual basis to incorporate a broader range of high-quality and newly published studies, ensuring alignment with evolving research trends and clinical standards. In parallel, to maintain quality, expand coverage, and ensure scalability, we plan to adopt a hybrid curation framework that combines expert manual review with AI-assisted text mining. Specifically, techniques such as named entity recognition, relation extraction, and automatic ontology mapping will be applied to screen new publications, highlight candidate associations, and assist curators in standardizing new entries. These enhancements aim to significantly improve scalability while maintaining high data quality.

Additionally, alongside ongoing technical enhancements, MTB-KB’s graph-based architecture and standardized entity integration strategy provide a flexible and transferable framework that can be readily applied beyond MTB. While the current focus is on MTB, the system is designed for seamless expansion to other high-priority pathogens, enabling the creation of multi-pathogen knowledgebases and supporting comparative analyses across infectious diseases. By demonstrating the feasibility of large-scale, literature-curated integration, MTB-KB establishes a methodological model for building comprehensive, evidence-driven resources across the broader landscape of infectious disease research.

In conclusion, MTB-KB provides a structured, literature-driven, and clinically relevant knowledgebase that complements existing TB resources. By combining manual curation, standardized annotation, and interactive visualization, it offers a practical and authoritative platform for researchers, clinicians, and public health professionals. Through the consolidation of influential findings across diverse domains of TB research, MTB-KB is well-positioned to strengthen global efforts in tuberculosis surveillance, control, and the development of evidence-based interventions, ultimately contributing to the WHO’s goal of ending the global TB epidemic by 2035.

## Data availability

MTB-KB is a curated knowledgebase of MTB-related studies at https://ngdc.cncb.ac.cn/mtbkb/.

## CRediT author statement

**Pan Li:** Data curation, Methodology, Software, Validation, Visualization, Writing – original draft, Writing – review & editing. **Cuidan Li:** Conceptualization, Methodology, Project administration, Validation, Writing – original draft, Writing – review & editing. **Rongxi Zhu:** Data curation, Validation. **Wenjing Sun:** Data curation, Validation. **Hengyu Zhou:** Software. **Zhuojing Fan:** Visualization. **Liya Yue:** Investigation. **Sijia Zhang:** Validation. **Xiaoyuan Jiang:** Investigation. **Quan Luo:** Conceptualization. **Jinying Han:** Data curation. **Hairong Huang:** Conceptualization. **Adong Shen:** Conceptualization. **Tuohetaerbaike Bahetibieke:** Conceptualization. **Jing Wang:** Conceptualization. **Wenbao Zhang:** Conceptualization. **Wen Hao:** Conceptualization. **Haitao Niu:** Conceptualization. **Congfan Bu:** Conceptualization. **Zhang Zhang:** Conceptualization. **Jingfa Xiao:** Conceptualization. **Renjun Gao:** Conceptualization, Writing – original draft, Writing – review & editing. **Fei Chen:** Conceptualization, Funding acquisition, Methodology, Project administration, Supervision, Writing – original draft, Writing – review & editing. All authors have read and approved the final manuscript.

## Competing interest

None declared.

## Acknowledgements

We thank a number of users for reporting bugs and providing suggestions. This work was supported by National Key Research and Development Program of China (Grant No. 2023YFC2604400, 2024YFC2309300, and 2024YFC2311300), and Open Research Project of the Key Laboratory of Viral Pathogenesis & Infection Prevention and Control of the Ministry of Education (Grant No. 2023VPPC-R09).

## Supplementary material

Supplementary material is available at *Genomics, Proteomics & Bioinformatics* online (https://doi.org/10.1093/gpbjnl/qzaxxxx).\

## Supplementary material

**Supplementary Figure 1 Flow diagram of literature selection**

The workflow for literature selection used to construct the knowledgebase. Publications were initially retrieved from the Web of Science.

**Supplementary Figure 2 Entity types and semantic relationship of the MTB-KB knowledge graph**

The ontology and semantic relationship framework of the MTB-KB knowledge graph. Entities are interconnected through predefined semantic relationships, including is_a (classification), reported_target_of (reported targeting relationship), has_effect_on (functional or biological effect), regulates (regulatory interaction), and contains (compositional relationship).

**Supplementary Table S1. Literature search queries for each MTB-KB research section**

**Supplementary Table S2. Controlled vocabulary used in MTB-KB**

## Notes

https://ngdc.cncb.ac.cn/mtbkb/

### Competing Interest Statement

The authors have declared no competing interest.

