## Supplemental Table 1 for "MTB-KB: A Curated Knowledgebase of *Mycobacterium tuberculosis* Related Studies"

**Supplementary Table S1. Literature search queries for each MTB-KB research section**

| Section | Query |
| --- | --- |
| Epidemiology | TS = (tuberculosis) AND TS = (outbreak OR epidemic OR prevalent OR transmission OR "contact tracing") |
| Diagnosis | TS = (tuberculosis) AND TS = (diagnosis OR diagnostic OR identification) |
| Treatment | TS = (tuberculosis) AND TS = (treatment OR therapy) |
| Vaccine | TS= (tuberculosis) AND TS= (vaccine) |
| Virulence factor | TS= (tuberculosis) AND TS= (virulence OR "virulence factor") |
| Drug resistance | TS= (tuberculosis) AND TS= ("drug resistance" OR "drug susceptibility" OR "drug sensitivity" OR "antibiotic resistance") |
| Immune response | TS= (tuberculosis) AND TS= (immune OR immunity) |
