## Supplemental Table 2 for "MTB-KB: A Curated Knowledgebase of *Mycobacterium tuberculosis* Related Studies"

**Supplementary Table S2. Controlled vocabulary used in MTB-KB**

| Data type | Description | Example |
| --- | --- | --- |
| Reported Features | Categorized biological or clinical themes identified within the article through keyword screening and manual review. | Drug |
| Experiment Type (resistance) | The methodology used to determine resistance. | Experimental Verification, Computer Analysis |
| Medium Type | Classification of the culture medium used. | Solid medium, Liquid medium |
| Virulence Factor Category | Broad functional classification of the virulence factor. | Immune modulation, Exotoxin, Nutritional/Metabolic factor, etc. |
| Change of Cell Frequency | The observed change in the proportion or frequency of the cell population. | Down, Up |
| Antibody Class | The isotype of the antibody being measured. | IgG |
| Trend | The direction of change in antibody titers or prevalence. | Upregulated, Downregulated |
| Experiment type | The classification of the experimental environment used to evaluate drug effect or resistance. | In vitro, In vivo, In silico |
| Resistance | Drug-resistance background of cases/isolates/cohort when applicable. | Drug-susceptible TB, RR-TB, MDR-TB, XDR-TB |
| WHO Category | WHO drug grouping defined in WHO TB treatment guidance. | Group A, Group B, Group C |
| Clinical Status | Clinical development/approval status of the drug. | Approved, Phase I, Preclinical, etc. |
| Route of Administration (Drug) | Route(s) of drug administration. | Oral, Intravenous. |
| Safety | Structured summary of drug safety/tolerability profile. | Well-tolerated, Intolerant |
| Vaccine Type | The biological platform or technology used to develop the vaccine. | Live-attenuated, Subunit, Viral vector |
| Vaccine Status | The current development phase or regulatory status of the vaccine. | Phase IIb, Phase III |
| Application Purpose | The intended clinical use of the vaccine. | Preventive, Therapeutic |
| Route of administration (Vaccine) | The physiological path by which the vaccine is introduced into the body. | Intradermal, Intramuscular |
| Effectiveness | A summary metric or qualitative assessment of the vaccine's protective efficacy. | Efficient, Inefficient |
