## Supplementary figures and images for "MTB-KB: A Curated Knowledgebase of *Mycobacterium tuberculosis* Related Studies"

### Supplemental Figure 1

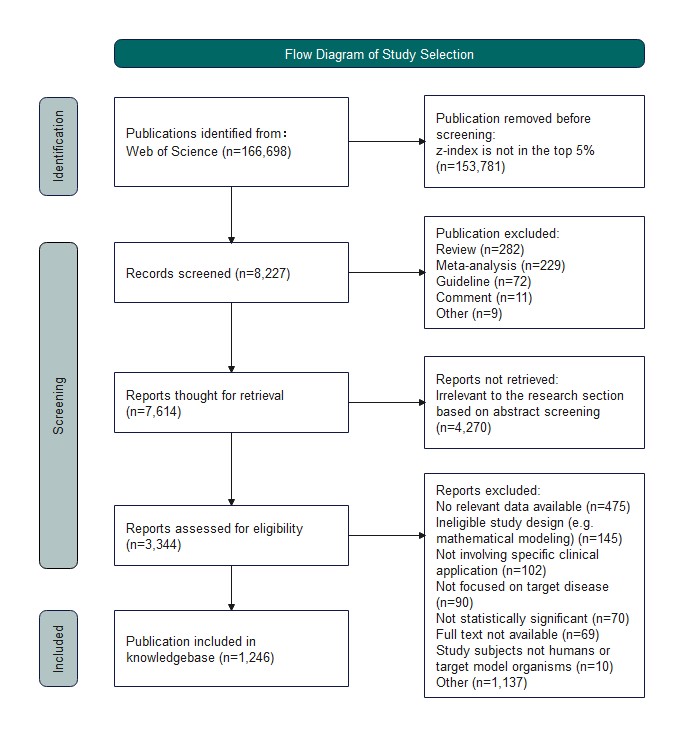

### Supplemental Figure 2

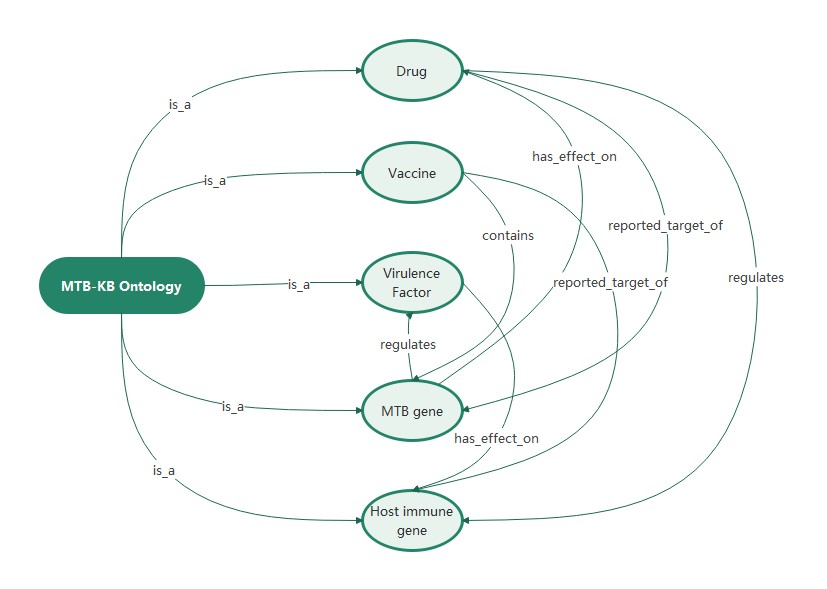
